# Comprehensive profiling reveals Sialyl-Tn upregulation and prognostic value in prostate cancer

**DOI:** 10.64898/2026.04.14.718221

**Authors:** Kirsty Hodgson, Libby Blencoe, Erin Smith, Aswini Sasikumar, Ziqian Peng, Margarita Orozco-Moreno, Richard Beatson, Paula A. Videira, Jennifer Munkley

**Affiliations:** Newcastle University Centre for Cancer, Newcastle University Institute of Biosciences, Newcastle, NE1 3BZ, UK; Comprehensive Cancer Centre, King’s College London. London. SE1 9RT; UCIBIO – Applied Molecular Biosciences Unit, Department of Life Sciences, NOVA School of Science and Technology | FCT NOVA, Universidade NOVA de Lisboa, Caparica, 2829□-□516, Portugal; Associate Laboratory i4HB - Institute for Health and Bioeconomy, NOVA School of Science and Technology | FCT NOVA, Universidade NOVA de Lisboa, Caparica, 2829□-□516, Portugal.; CDG & Allies – Professionals and Patient Associations International Network (CDG & Allies – PPAIN), Caparica, 2829□-□516, Portugal

**Author notes:** Correspondence to: Paula A. Videira or Jennifer Munkley.

## Abstract

Prostate cancer is a common cancer in males and there is an urgent unmet clinical need to identify new therapies for advanced disease. Aberrant glycosylation is common in prostate cancer and plays a functional role in disease progression. The sialyl-Tn antigen (sTn) has been widely studied in cancer, yet its involvement in prostate cancer remains relatively unexplored. Here, we utilise a novel anti-sTn antibody (L2A5) to comprehensively monitor sTn expression levels in clinical prostate cancer tissues encompassing normal, benign, primary, metastatic castrate-resistant prostate cancer (CRPC), and patient-derived xenografts (PDXs). We show that while sTn is detected at low or negligible levels in normal prostate tissues, it is expressed in 44% of prostate tumours, and prostate cancer patients with high sTn levels have significantly poorer survival times. Analysis of metastatic therapy resistant prostate-derived tumours growing in liver and bone, shows sTn is expressed in 37.5% of cases. Furthermore, we show sTn is expressed in nearly half of PDXs tested, supporting the use of PDX models as tools for testing anti-sTn therapeutic strategies. These findings identify sTn as potential prognostic biomarker and therapeutic target in prostate cancer and lay the groundwork for the development of sTn-targeted precision therapies for advanced disease.

## Introduction

Prostate cancer is a common cancer in males leading to more than 350,000 deaths worldwide every year (1). Androgens are required for normal prostate development and function, however in prostate cancer the androgen receptor (AR) signalling axis is hijacked to promote disease progression (2). Drugs that block the production of androgens and/or inhibit AR activity are the cornerstone treatment for advanced prostate cancer, and new AR targeted therapies have improved patient outcomes (3-5). However, patients still inevitably develop resistance to hormonal therapy, known as castrate resistant prostate cancer (CRPC) (6), which remains lethal with a median overall survival ranging from 2 to 3 years (7, 8). Additional therapeutic strategies are urgently needed to improve outcomes for prostate cancer patients with advanced prostate cancer.

Aberrant glycosylation is a hallmark of cancer (9), and in prostate cancer changes to glycans have been functionally linked to disease progression (10). *O*-glycans are found on ∼80% of proteins travelling through the secretory pathway and represent one of the most abundant and diverse glycan types [8]. In tumour cells, the processing of *O*-glycans into branched structures can be disrupted, leading to the expression of the cancer-associated sialyl-Tn (sTn) antigen, which is a truncated *O*-glycan containing a sialic acid α-2,6 linked to GalNAc on serine or threonine residues on glycoproteins. sTn has negligible expression in healthy tissues but is detected on many cancers, such as bladder, ovarian, colon, breast and pancreatic cancers (11-18), where it is associated with poor patient prognosis (19, 20). Mechanistically, sTn plays a role in promoting tumour progression, metastasis, immune evasion, and therapy resistance, and therefore represents an attractive target for the development of anti-cancer strategies (21, 22). Over the years, numerous anti-sTn monoclonal antibodies have been developed, including B72.3, HB-Stn1, CC49, TKH2, and 3F1 with varying fine specificities (14, 23-28). However, anti-glycan monoclonal antibodies often have low affinity, poor selectivity, and mixed specificity (28-30). Recently, a novel monoclonal antibody, known as L2A5 was developed using hybridoma technology that is highly specific to the sTn antigen (31). L2A5 shows a unique binding pattern specific to invasive regions and metastatic sites in tumours that other anti-sTn antibodies fail to detect (31, 32). Notably, L2A5 exhibits fine specificity towards cancer-associated MUC1 and MUC4 mucin derived glycopeptides (32), potentially enabling selective targeting of tumour cells over healthy cells beyond the recognition of the sTn antigen alone (31). Preclinically, L2A5 also showed therapeutic potential as chimeric antigen receptor against models of triple negative breast cancer (TNBC) (33, 34). In TNBC patient tissues, L2A5 binds with 23.8% of cases and correlates with significantly reduced survival times in patients, lower c-Myc expression, and an immunosuppressive tumour microenvironment (17).

In prostate cancer, historical data suggests sTn is expressed in up to 80% of high-grade prostate tumours (26, 35, 36). However, these studies were carried out more than 30 years ago, involved limited clinical samples, and utilised anti-sTn antibodies with now-recognised limitations of specificity and cross-reactivity (14, 23-28). To date, a comprehensive analysis of sTn across the full clinical spectrum of prostate tumours has not yet been reported and remains a critical research gap in the field (37). Here, we address this knowledge gap, by studying sTn expression in normal prostate tissues, benign prostate hyperplasia (BPH) tissues, primary prostate tumours from both White and Black patients, in metastatic CRPC tumour tissues, and in PDX samples. Recognising the limitations of older anti-sTn antibodies, we chose to utilise the newly developed L2A5 anti-sTn antibody, which is validated to outperform commonly used anti-sTn antibodies (31, 32). Our findings show sTn expression is upregulated in prostate cancer, with sTn specifically detected in prostate tumour cells, and negligible levels detected in normal prostate tissues. We find that high levels of sTn correlate with significantly reduced survival times in prostate cancer patients, and that sTn is expressed in 37.5% of therapy resistant metastatic prostate tumours, including prostate-derived cancers that have spread to liver and bone. Furthermore, sTn is expressed in prostate cancer patient-derived xenografts (PDXs), identifying a potential tool for evaluating therapeutic approaches to target sTn for prostate cancer.

## Methods

### L2A5 antibody

The monoclonal antibody L2A5 is a proprietary anti-sTn monoclonal antibody and has been previously described (PCT: WO2019147152A1) (31).

### Clinical samples

#### TMA cohort 1

A 96 case TMA comprising cores of normal prostate tissue and prostate adenocarcinoma of different Gleason grades was purchased from US Biomax (PRC1921-L38).

#### TMA cohort 2

A 100 case prostate cancer and benign prostate tissue TMA with survival data was purchased from US Biomax (MPR1005sur) (38).

#### TMA cohort 3

The CHTN_PrC_Prog1 TMA, containing prostate tumour samples from 7 Black and 23 White prostate cancer patients, has been previously published (39) and was obtained through Cooperative Human Tissue Network (CHTN) at the University of Virginia.

### Metastasis tissue samples from rapid autopsy (tissue cohort 4)

40 cases of rapid autopsy FFPE tissue samples from prostate-derived tumours growing in bone or liver were kindly provided by Dr Colm Morrissey (University of Washington) via the Prostate Cancer Biorepository Network (PCBN). Biopsies of metastatic sites were obtained from patients with CRPC within hours of death using a cordless drilling trephine (DeWalt Industrial Tool) and model 2422-51-000 trephine (DePuy). Liver cores were fixed in 10% neutral buffered formalin and paraffin embedded. Bone cores were fixed in 10% neutral buffered formalin, decalcified with 10% formic acid and paraffin embedded.

### LuCaP Patient-derived xenograft (PDX) TMA

The LuCaP PDX TMA is previously published and contains 39 CRPC PDX model tumours, established from distinct patients from specimens acquired at either radical prostatectomy or at autopsy, implanted and grown subcutaneously (in triplicate) in intact or castrated mice (40, 41).

### Immunohistochemistry

IHC was performed with the IHC Prep & Detect Kit for Rabbit/Mouse Primary Antibody (Proteintech, PK10019) following the manufacturer’s protocol with the following exceptions. Slides were dewaxed in Histo-Clear (SLS, NAT1330) and antigen retrieval was performed with Sodium Citrate antigen retrieval buffer (Proteintech, PR30001). Slides were incubated with 2.5 µg/mL anti-STn monoclonal antibody L2A5 overnight at 4°C. Coverslips were mounted with Histo-Mount (SLS, NAT1308) and images were acquired with the ZEISS Axioscan 7 Slide Scanner at 20X magnification. Images were processed using OMERO Plus (Glencoe Software). The TMAs shown in Figures 1-3 were scored by a pathologist using the 0–300 HistoScore method as described previously (38, 42, 43). Only epithelial cells were scored. For the PDX TMA shown in Figure 5, images were analysed using QuPath (version 0.6) software (each PDX sample was analysed in triplicate samples from the same mouse, and across three independent mice). Positive cells were outlined and classified as having low, moderate or high staining intensity to calculate a 0-300 HistoScore, calculated as the sum of each staining intensity score (1+, 2+, 3+) multiplied by the percentage of cells classified at that intensity using the equation staining Index (H-score) = Σ (each intensity score × % of cells at that intensity).

**Figure 1.**
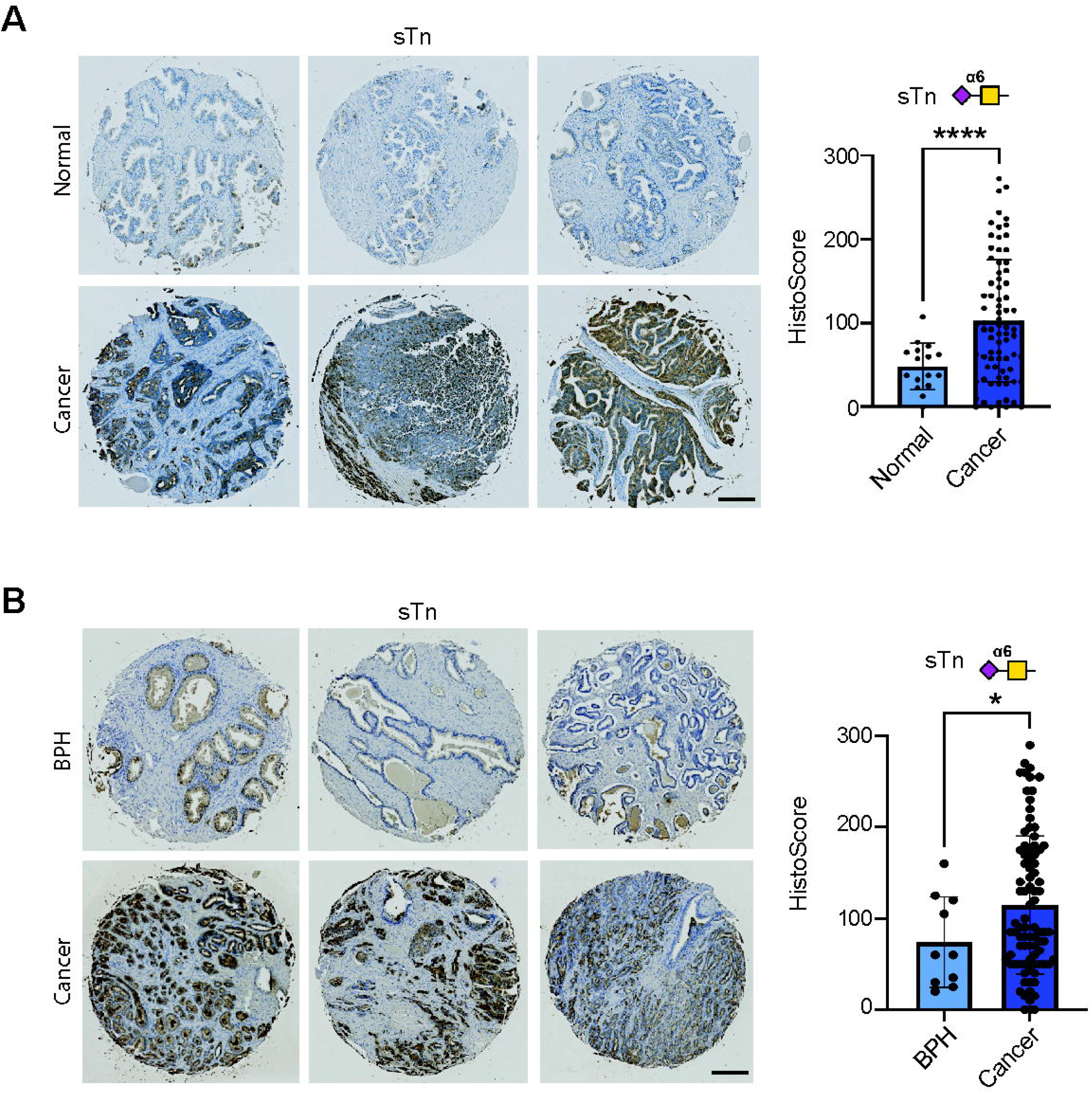
The sTn antigen is expressed in prostate tumours. (**A**) Analysis of sTn using L2A5 immunohistochemistry in a 96-case tissue microarray (TMA) comprising 17 normal prostate tissue samples and 79 samples of prostate tumour tissue (TMA cohort 1) showed that sTn is detected at low or negligible levels in normal prostate tissue and is significantly upregulated in prostate tumour tissues (unpaired t test, p<0.0001). Furthermore, sTn was detected at high levels (Histoscore >100) in 44% of prostate cancer cases. (**B**) L2A5 immunohistochemistry analysis of sTn expression in a previously published 100 case TMA (38) containing 10 benign prostate hyperplasia (BPH) tissue samples and 90 prostate cancer tissue samples (TMA cohort 2). sTn was detected at significantly higher levels in prostate cancer tissues compared to BPH tissue (unpaired t test, p=0.0156), with sTn detected at high levels (Histoscore >100) in 46% of prostate cancer cases. Scale bar is 200 µm.

## Statistical analysis

Statistical analyses were conducted using the GraphPad Prism software (version Prism 10) using suitable tests as described in the legends. Data are presented as the mean of three independent samples ± standard error of the mean (SEM). Statistical significance is denoted as *p<0.05, **p<0.01, ***p<0.001 and ****p<0.0001.

### Study approval

All human studies were reviewed by appropriate ethics committee and performed in accordance with the ethical standards laid down in the Declaration of Helsinki. All autopsy tissues were collected from patients who had signed written informed consent under the aegis of the Prostate Cancer Donor Program at the University of Washington (41). The IRB of the University of Washington approved this study. All patient-derived xenograft experiments were approved by the University of Washington IACUC. For animal experiments, the ‘Principles of Laboratory Animal Care’ were followed as well as specific national laws as detailed in previous publications (40, 41).

## Results

### 1. The cancer-associated sTn antigen is upregulated in prostate tumours

Previous studies suggest that the sTn antigen is expressed in prostate cancer tissues (26, 35, 36). To overcome the limitations of the anti-sTn antibodies used in previous studies, we utilised the L2A5 anti-sTn monoclonal antibody, which has stood out for its reported specificity towards sTn expressing tumours tissues (31). Using L2A5 immunohistochemistry, we monitored the levels of sTn in three distinct patient cohorts. In line with previous studies, only epithelial cells were scored, and a Histoscore <50 was considered negative, scores of 50-100 were weakly positive/negligible, and scores >100 were recorded as positive (44, 45). In a 96-case tissue microarray (TMA) containing 17 normal prostate tissue samples and 79 samples of prostate tumour tissue, sTn was significantly higher in prostate tumours tissues compared to normal prostate tissues (p<0.001) (**Figure 1A**), and was expressed in 44% prostate of cancer tissues tested (sTn Histoscore >100). sTn was absent in 28% of prostate cancer tissues (sTn Histoscore <50), with a further 28% of cases having negligible levels of sTn (sTn Histoscore 50-100). sTn was absent or detected negligible levels in 100% of normal prostate tissues (sTn Histoscore <100). Further analysis of a 100 case TMA, containing 10 benign prostate hyperplasia (BPH) tissue samples and 90 prostate cancer tissues, revealed sTn was highly expressed in 46% of prostate tumours (sTn Histoscore >100), with levels significantly higher in prostate tumours compared to BPH samples (p=0.0156) (**Figure 1B**). Consistent with with previous studies in breast cancer tissues, all sTn positive cases in our cohorts showed membrane staining. However, some cases also exhibited additional cytoplasmic signal, which is likely a reflection of the biosynthetic pathway of sTn (17). Our findings, using a well validated highly specific antibody, show the cancer-associated sTn antigen is significantly upregulated in prostate tumours compared to normal or benign prostate tissues.

### 2. Expression of sTn in Black prostate cancer

The data presented above reveals sTn is expressed in prostate tumour tissues. As Black patients with prostate cancer have higher incidence rates, worse outcomes, and differences in tumour biology (46), we next investigated if the levels of the sTn antigen might be altered in Black prostate cancer. Analysis of sTn levels in a TMA comprising 7 Black prostate cancer tissues and 23 White prostate cancer tissues revealed sTn is expressed (sTn HistoScore >100) in 6/6 (100%) of Black tumour tissues and 20/24 (83%) of White tumour tissues Overall, we detected no significant difference in sTn levels between Black and White prostate cancer (n=30, unpaired t test, p=0.4603) (**Figure 2**). Taken together, these findings show the sTn antigen is expressed at similar levels in White and Black prostate tumour tissues, and that once anti-sTn therapies become available for cancer therapy they will likely be relevant to both White and Black prostate cancer patients.

**Figure 2.**
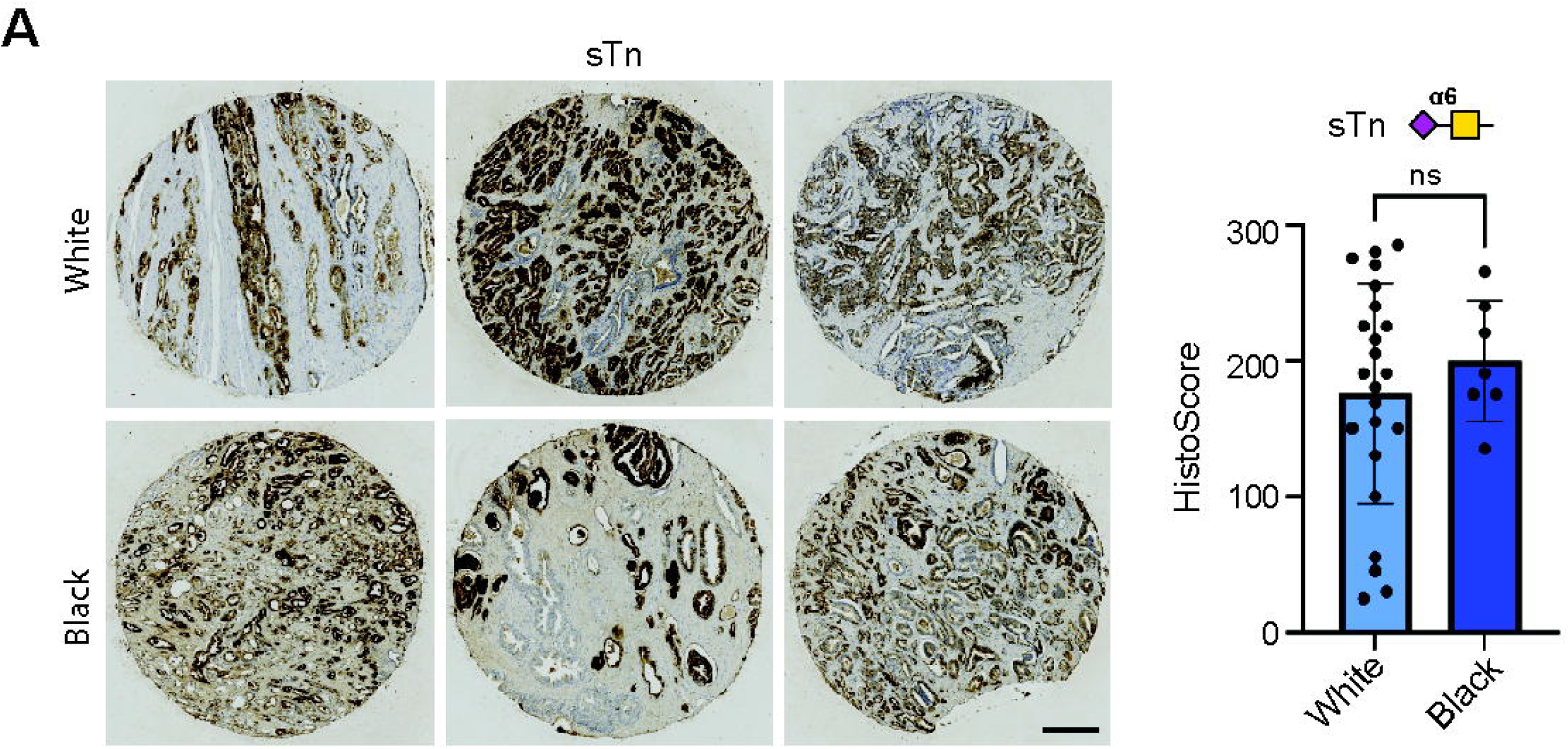
sTn is expressed in Black prostate cancer. L2A5 immunohistochemistry analysis of a previously published TMA (TMA cohort 3) (39) comprising prostate tumour samples from 7 Black patients and 23 White patients (TMA cohort 3). No significant differences in sTn levels were detected in prostate tumours from White and Black patients (n=30, unpaired t test, p=0.4603).

### 3. High sTn expression correlates with reduced survival times in prostate cancer patients

Next, to investigate the prognostic and therapeutic potential of sTn in prostate cancer, we further analysed the association between sTn expression and clinical parameters. No correlation was found between sTn levels and Gleason grade across two independent patient cohorts (TMA cohorts 1&2) (Supplementary Figure 1). As sTn correlates with poor patient prognosis in breast cancer (17), we next tested if expression of sTn might also correlate with survival times in prostate cancer patients. We analysed a previously published TMA (TMA cohort 2) comprising 90 cases of primary prostate cancer tissues (47). Here, when we stratified prostate cancer patients based on high and low sTn levels (defined as the top and bottom 25^th^ percentile of expression) patients with high sTn levels had significantly reduced survival rates compared to patients with low sTn levels (p=0.0319 (**Figure 3**). Our data shows high sTn levels correlate with poorer patient prognosis in prostate cancer and provides rationale for future studies investigating anti-sTn therapeutic strategies for prostate cancer.

**Figure 3.**
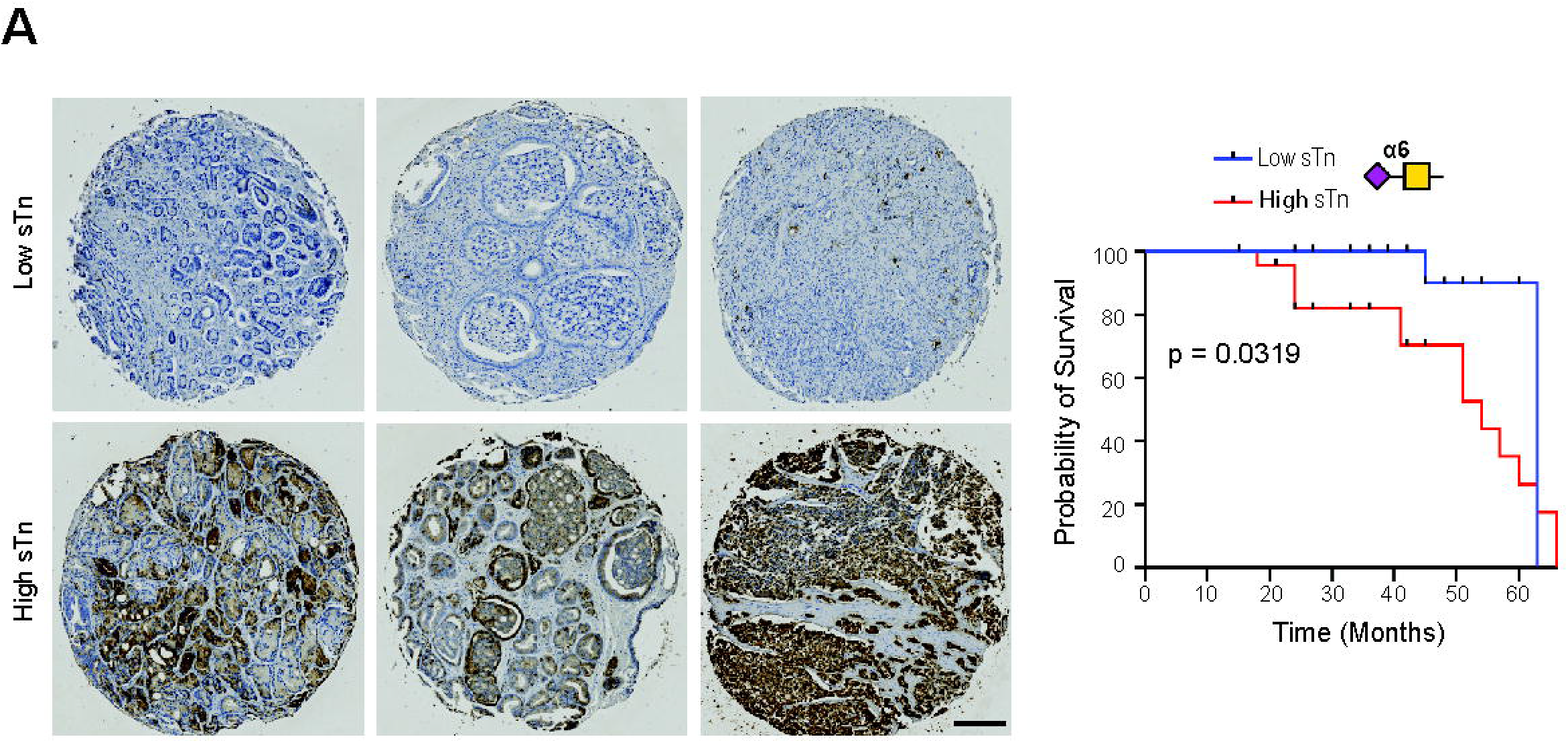
sTn levels correlate with poor patient prognosis in prostate cancer patients. Additional analysis of sTn expression in a 100-case prostate cancer TMA using L2A5 immunohistochemistry (TMA cohort 2). Stratification of patients based on high and low sTn Histoscore levels shows patients with high sTn levels (defined as the top 25^th^ percentile of expression) had significantly poorer survival rates compared to patients with low sTn levels (defined as the bottom 25^th^ percentile of expression) (n=100, Kaplan-Meier regression model, p=0.0319) Scale bar is 200 µm.

### 4. sTn is expressed in metastatic therapy resistant prostate cancer

Nearly all men with advanced prostate cancer who receive hormone therapy eventually progress to CRPC, where metastasis is common (48). Although previous studies have analysed the levels of glycosyltranferase enzymes and changes to *N*-glycans in CRPC (49-51), the expression levels of sTn in metastatic therapy resistant disease has not yet been evaluated. To address this research gap, we next monitored sTn in 40 rapid autopsy tissues obtained from lethal prostate-derived liver and bone metastatic tumours. L2A5 immunohistochemistry showed sTn was detected in 15/40 CRPC samples tested, including 8/20 bone tumours (**Figure 4A**) and 7/20 liver tumour tissues (**Figure 4B**). Our findings show sTn is highly expressed in lethal metastatic prostate tumours and identify the sTn antigen as potential therapeutic target for the development of new therapies for advanced disease.

**Figure 4.**
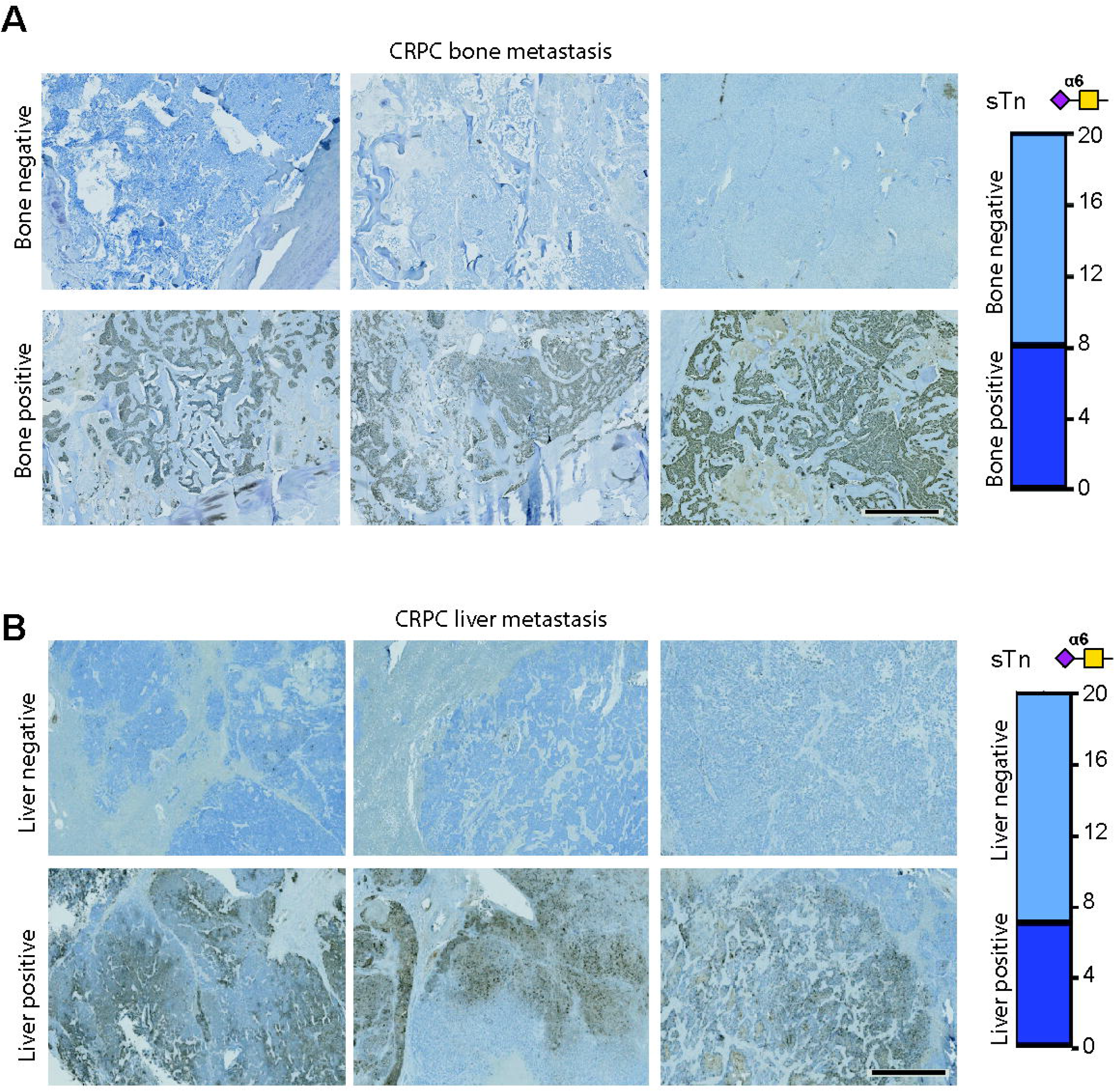
sTn is expressed in lethal metastatic therapy resistant prostate cancer. Analysis of sTn levels in 40 metastatic CRPC tumours obtained via rapid autopsy (tissue cohort 4). (**A**) L2A5 immunohistochemistry shows sTn is expressed by 8/20 prostate-derived tumours growing in bone, and (**B**) 7/20 prostate-derived tumours growing in soft tissue (liver). Scale bar is 2000 µm.

### 5. sTn expression validation in prostate cancer patient-derived xenografts (PDXs)

Next, with the longer-term goal of identifying a clinically relevant platform for the future evaluation of anti-sTn therapeutic for prostate cancer, we assessed sTn expression within the LuCaP prostate cancer PDX series, which reflects the diverse molecular composition of human CRPC (40, 41). Using L2A5 immunohistochemistry, we evaluated 39 PDXs, representing a spectrum of metastatic sites and histologies, grown in both intact and castrated immunodeficient mice. Among the 39 PDXs evaluated (each established from distinct patients), after grafting patient tumour cells into a new microenvironment, 19 showed positive sTn expression and 20 PDXs tested showed no sTn expression. Some sTn-positive samples showed moderate sTn staining, while others displayed strong positive staining for sTn, with sTn-positive staining detected in PDXs established from ascites, and from liver and bone metastatic tissues (**Figure 5**). Overall, there was no significant difference between PDX samples grown within intact or castrated mice (**Figure 5A**). However, LuCaP 147 demonstrated a significant increase in sTn expression in castrate conditions (**Figure 5B**) and, LuCaP 136 displayed a significant decrease in sTn expression when grown in castrated mice (**Figure 5C**). Interestingly, in matched PDX tissues from the same patient (LuCaP 189), sTn levels were significantly increased in PDX models established from a bone tumour compared to from an adrenal tumour (n=6, paired t test, p=0.0168) (**Figure 5D**). Taken together, our data shows the LuCaP PDX models retain sTn expression as a key feature of patient tumours, suggesting these models have the potential to accelerate anti-sTn drug development.

**Figure 5.**
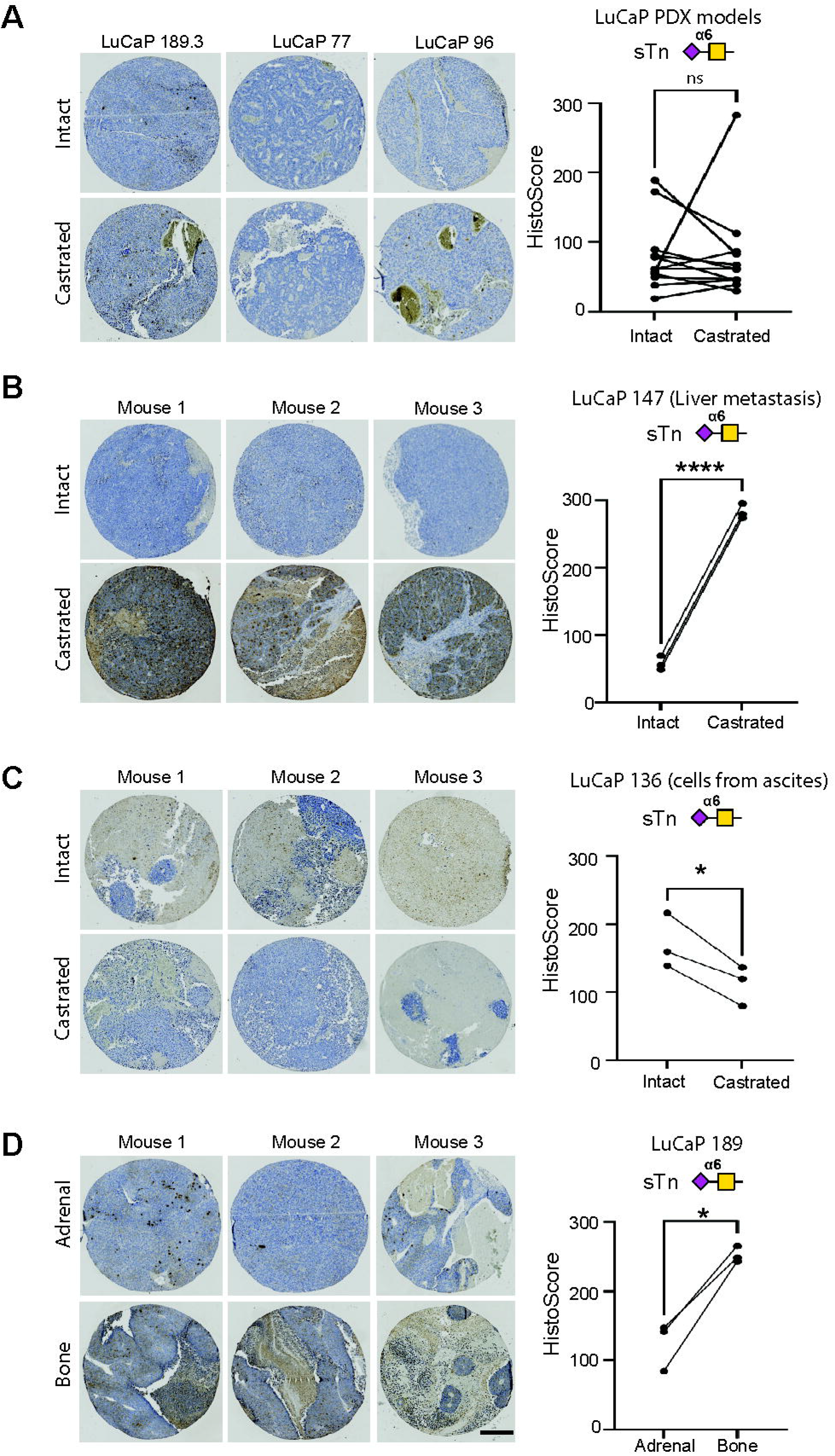
Analysis of sTn in LuCaP patient-derived xenografts (PDX) tissues. L2A5 immunohistochemistry was used to analyse the sTn antigen in tissue samples from the LuCaP PDX series, comprising PDX samples from 39 patients with CRPC grown in intact immunocompromised mice (12 with matching samples grown in castrated mice) (40). (**A**) Analysis of sTn Histoscores showed no significant difference in sTn levels in 12 PDX matching samples grown in intact or castrated conditions (paired t test, n=0.9693). Representative images for 3 PDX models are shown. Each PDX sample was analysed in triplicate within 3 mice. (**B**) In LuCaP 147, which was established from implanted tissue from liver metastasis, the levels of sTn are significantly increased in PDX samples grown in castrated conditions compared to in intact mice (n=6, paired t test, p<0.0001). (**C**) For LuCaP 136 (grown from cells from ascites), there was a significant reduction in sTn levels in PDX tissues grown in castrated mice compared to intact mice (n=6, paired t test, p=0.0366). (**D**) Comparison of sTn levels in PDX tissues established from adrenal metastasis (LuCaP 189.3) and bone metastasis tissue (LuCaP 189.4) from the same patient shows sTn is significantly increased in PDX samples established from bone metastatic tissue (n=6, paired t test, p=0.0168). Representative images are shown. Scale bar is 200 µm.

## Discussion

The truncated *O*-glycan structure sTn has emerged as a compelling target for cancer therapeutics due to its aberrant yet specific expression pattern on epithelial tumours. While detection of sTn is rare or absent in normal tissues, sTn is prominently expressed in various epithelial tumours, and targeting sTn holds tremendous potential to treat a wide range of solid tumours. In prostate cancer, historical studies have detected sTn in up to 50-80% of prostate tumours (25, 30, 31). However, these studies were based on limited sample sizes, and utilised benchmark antibodies which now have recognised limitations. For example, B72.3 shows a clear preference for serine over threonine residues (52, 53), and retains around 26% of binding to sTn positive cells treated with sialidase (31). Given the major importance held by the accurate detection of sTn in prostate cancer for advancing the development of targeted therapies, we utilised the L2A5 anti-sTn antibody, which overcomes key limitations of previous anti-sTn antibodies, to monitor the sTn antigen in prostate tumour tissues. Our study provides a novel and comprehensive characterisation of the sTn antigen in prostate cancer tissues representing the full clinical spectrum of the disease. We report 44% of prostate tumours express sTn with selective staining of malignant tissues and limited reactivity toward normal prostate tissue. Consistent with previous studies in other cancer types (17, 54, 55), high sTn levels correlate with significantly poorer survival outcomes in prostate cancer patients. Furthermore, we show sTn remains expressed in lethal therapy resistant prostate tumours obtained via rapid autopsy and in clinically relevant CRPC PDX models. Our findings underscore the promise and provide rationale for future pre-clinical studies exploring the development of sTn-targeted therapies or prostate cancer.

Previous studies have suggested that both ST6 N-acetylgalactosaminide alpha-2,6-sialyltransferase 1 (ST6GalNAc1), the main enzyme responsible for the synthesis of sTn (56), and sTn itself can be regulated by androgens in prostate cancer cells (57, 58). The findings presented in this study show that expression of sTn may is likely context dependent and may depend on the specific tumour microenvironment, highlighting the complex nature and molecular landscape of prostate cancer biology. The sTn antigen has a well-established role in promoting immune evasion and contributing to an immunosuppressive tumour microenvironment (22, 59). Studies show sTn can bind immune cells via specific lectins, including Sialic acid-binding immunoglobulin-type lectins (Siglecs) and Macrophage Galactose-type Lectin (MGL) to promote tumour immune evasion (60, 61). In bladder cancer, sTn-expressing cancer cells impair dendritic cell maturation and suppress anti-tumour T cell responses (22), and in breast cancer sTn expression is linked to M2 polarization (17). Although the specific functional role of sTn in prostate tumour immune suppression is a research knowledge gap, recent findings show Siglec-7, 9, -10 and 15, which can engage sTn (21, 62-65) are expressed by immunosuppressive macrophages and NK cells in the prostate tumour immune microenvironment (38, 66, 67). However, although tumour-associated macrophages are heavily infiltrated into prostate tissues (68), whether these express MGL is yet to be reported. Together, these findings underscore the likely significant role of the sTn antigen in promoting prostate tumour immune suppression which correlates to poorer disease outcomes, and points to the need for further studies investigating how both the Siglec-sTn and the MGL-sTn axis might contribute to prostate cancer progression.

The expression of sTn on the surface of cancer cells makes it a promising target for cancer therapy, particularly as healthy adult cells do not express sTn, which will prevent off target toxicity. The Theratope vaccine, which targets sTn, was evaluated in Phase I and II clinical trials, and succeeded in triggering T cell-dependent responses against cancer cells in breast and ovarian cancer patients (69), but failed to show improved overall survival in phase III clinical trials (70); potentially as the trial did not stratify/select sTn+ patients for inclusion which later studies indicate is important (71). Additional strategies being explored to block or target sTn include CAR and antibody-based approaches. Antibodies targeting sTn have emerged with different specificities against the epitope (27, 31, 53, 72, 73). Targeted therapy based on antibodies is a rapidly expanding field, with >150 monoclonal antibodies receiving FDA approval in the last 40 years (74), and antibodies targeting sTn hold promise to enhance immune responses against sTn positive tumours. The monoclonal antibody 3P9 has been reported to directly inhibit the growth of colorectal tumours (73), and an sTn antibody-drug conjugate (ADC) has demonstrated efficacy against sTn-expressing tumours in vivo (25). The clinical potential of anti-sTn antibody-drug conjugates is demonstrated by SGN-STNV, which was evaluated in a clinical trial for solid tumours (NCT04665921). Furthermore, anti-STn CAR-T cell therapies have shown significant anti-cancer activity in pre-clinical mouse models (33, 34, 53), and anti-TAG72 antibodies that target mucin enriched sTn are being investigated in clinical trial for the treatment of ovarian cancer (NCT05225363).

In conclusion, here we present of comprehensive analysis of the cancer-associated sTn antigen in prostate cancer tissues representing both untreated primary disease as well as therapy resistant metastatic tumours. Using a novel anti-sTn antibody (L2A5) with unique binding specificity to sTn, we show the sTn antigen is upregulated in prostate tumour tissue and correlates with poorer survival rates in patients. Furthermore, sTn remains expressed in metastatic CRPC. Our findings introduce the sTn glycan epitope as a promising therapeutic target in prostate cancer and identifies the LuCaP series of PDXs (40) as a pre-clinical platform for the future testing of anti-sTn agents.

---

**Supplementary Figure 1.**
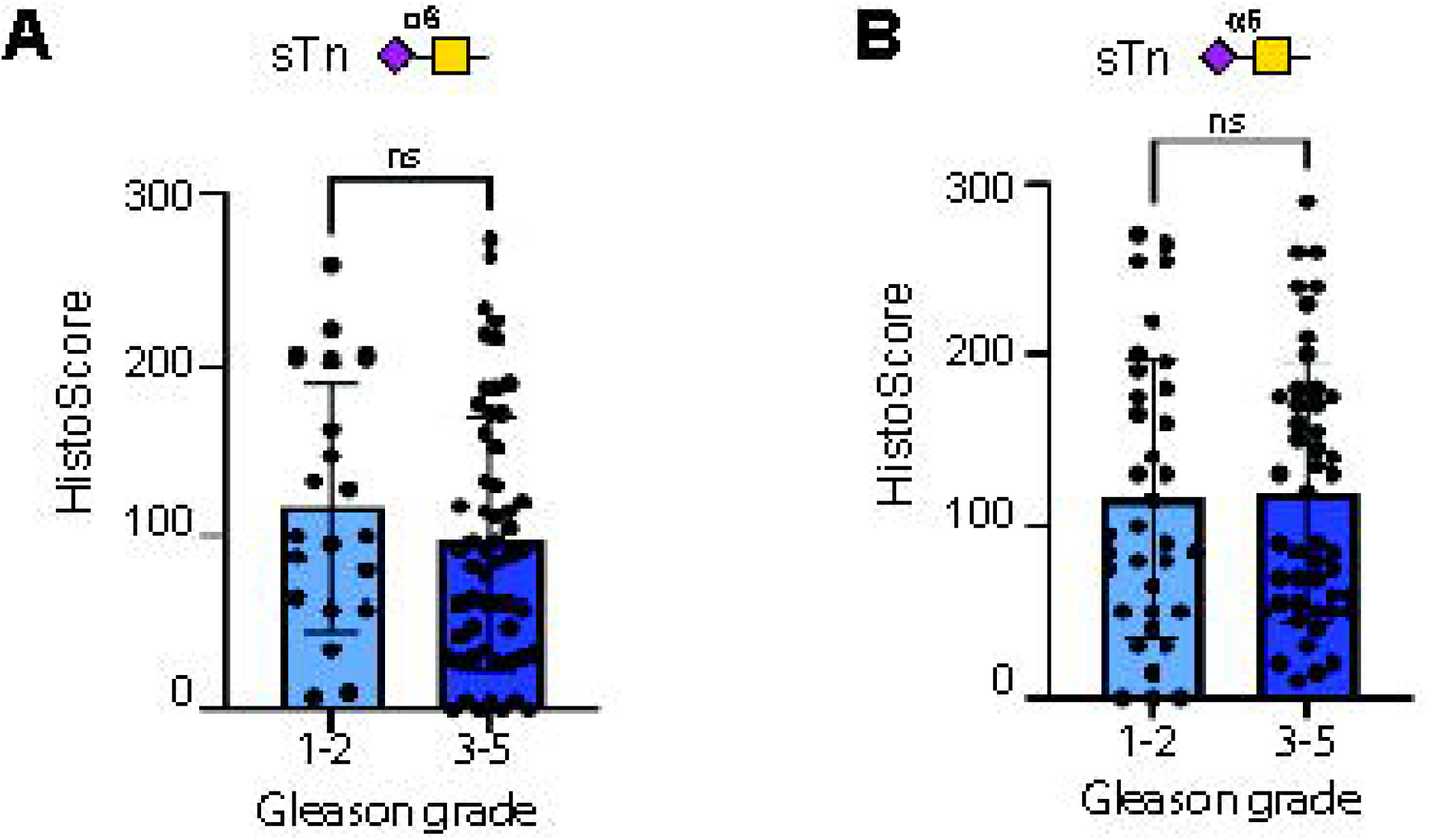
Expression of sTn does not correlate with prostate cancer Gleason grade. Analysis of sTn levels in TMA cohort 1 and 2 and correlation with Gleason grade. (**A**) Analysis of sTn levels in prostate cancer tissue samples from TMA cohort 1 shows there is no significant difference in sTn levels in Gleason grade 1-2 tumours compared to Gleason grade 3-5 tumours (n=79, unpaired t test, p=0.2820). (**B**) Analysis of sTn levels in 90 prostate cancer tissue samples from TMA cohort 2 shows there is no significant difference in sTn levels in Gleason grade 1-2 tumours compared to Gleason grade 3-5 tumours (n=90, unpaired t test, p=0.8917).

## Acknowledgements

We thank the patients and their families, Pete Nelson, Evan Yu, Heather Cheng, Bruce Montgomery, Jessica Hawley, Mike Schweizer, Daniel Lin, Funda Vakar-Lopez, Michael Haffner, Martine Roudier, Lawrence True, Meagan Chambers, Colm Morrissey and the rapid autopsy teams for their contributions to the University of Washington Medical Center Prostate Cancer Donor Rapid Autopsy Program.

## Ethics approval

The prostate cancer tissue samples analysed in Figures 3 and 4 were kindly provided by the Prostate Cancer Biorepository Network (PCBN). Written informed consent was obtained from all patients. The PCBN ethics committee reviewed our project and provided ethical approval for use of the samples in our project (refs: 191112.1, 210412.1 and 210203.1).

